# Impact of maternal obesogenic diet on maternal and offspring microbiome development

**DOI:** 10.1101/2024.01.21.576569

**Authors:** Kyoko Hasebe, Michael D Kendig, Nadeem O Kaakoush, Aynaz Tajaddini, R Frederick Westbrook, Margaret J Morris

## Abstract

Pregnancy can alter gut microbiota composition, but how an obesogenic diet impacts maternal gut microbiota, and the extent to which this influences offspring microbiome can be obscured by confounding factors. This study examined changes in gut microbiota composition across pre-pregnancy, gestation and lactation in rat dams fed either a high-fat, high-sugar Cafeteria (Caf) diet or Chow. Microbiome development was assessed in their offspring weaned onto chow. Caf diet consumption during pregnancy increased weight gain and adiposity, and compromised subsequent maternal nursing behaviour. α- and β diversity measures in Caf-fed dams showed a different trajectory across the progression of pregnancy, with no change in *Bacteroidetes* and *Firmicutes* abundance compared with Chow dams. Offspring born to Caf dams exhibited greater adiposity and plasma leptin at weaning and 14 weeks of age than those born to Chow dams. Maternal Caf diet induced clear differences in β diversity in weanlings but not α diversity. SourceTracker analysis revealed similarities in the gut microbiota of Chow weanlings and maternal gut microbiota in lactation, whereas the microbiota of Caf weanlings was similar to the maternal gut microbiota during gestation. Maternal Caf diet exerted only marginal effects on gut microbiota composition in 14-week-old offspring.

## Introduction

During pregnancy, physiological changes to the immune and endocrine systems nourish the developing fetus and impact metabolic status. While current evidence suggests that gut microbiota composition is strongly associated with physiological changes during pregnancy (Abu-Raya et al., 2020; Tian et al., 2023), observations regarding the temporal dynamics of any changes in the maternal gut microbiome during pregnancy are inconsistent. In humans, Koren et al. (2012) reported changes in maternal gut microbiota diversity from the first to third trimester, specifically increases of *Proteobacteria* and *Actinobacteria*, and decreased α diversity. In contrast, DiGiulio et al. (2015) found that pregnancy progression was not associated with α-and β diversity in weekly samples from vagina, stool, saliva and tooth/gum. Similarly, Yang et al. (2020) found limited gestational age-associated variations in human gut microbiota, while noting variability associated with host factors such as age, pre-pregnancy body mass index (BMI) and disease states. Finally, Rasmussen et al. (2020) reported a significant increase in Shannon diversity, but no difference in Faith’s Phylogenetic Diversity, from weeks 24 to 36 of pregnancy, a decline in *Lactobacillus* relative abundance from week 24 until birth, and no detectable change in β diversity.

Fluctuations in maternal microbiota have also been observed over gestation in animal models. Changes in relative abundances of *Clostridium* spp., *Akkermansia muciniphila*, *Methanobrevibacter* spp., *Bacteroides/Prevotella* spp., and *Roseburia* spp. were observed in faecal samples from pregnant Sprague-Dawley rats from pre-pregnancy, gestation day 14 and lactation day 19 (Paul et al., 2018). Using a porcine model, Ji et al. reported increased microbiota α diversity and differences in β diversity across gestation and lactation (2019). Zhang et al. (2023) reported increases in two α diversity indices (Shannon and Faith’s Phylogenetic Diversity) during gestation in goats compared with baseline, while differences in maternal microbiota β diversity from baseline to gestation and lactation were driven by decreased relative abundance of *Anaerovoracaceae* Family XIII AD3011 group at gestation. They also reported that alterations in pro- and anti-inflammatory cytokine levels in serum were accompanied by changes in maternal microbiota. The above noted differences between human and animal data regarding changes in maternal gut microbiota across pregnancy and lactation could relate to the inability to precisely control diet and other potential confounding factors in human microbiota studies during pregnancy.

Maternal microbiota is transferred to offspring through various routes. While adult gut microbiota is dominated by *Firmicutes* and *Bacteroidetes* phyla, *Proteobacteria* and *Actinobacteria* are predominant in early life, when the gut microbiota is highly unstable and variable (Milani et al., 2017). However, once established, the microbial community is stable and can fully recover even after multiple antibiotic exposures (Lavelle et al., 2019; Rashidi et al., 2021). Hence, early life gut microbiota serves as a foundation of the gut microbial community in later life. However, little is known regarding how any changes in maternal gut microbiota during pregnancy contribute to infant gut microbiota, which is known to be affected by multiple factors, a key one being maternal diet. Using a primate model, Ma et al (2014) tested the effects of a maternal high fat (HFD) or control diet on the early life offspring gut microbiome. They found that maternal diet during the perinatal period, not maternal obesity *per se*, appeared to shape offspring gut microbial community in the first year of life. In a human study of 163 mother-child dyads in which mothers were identified as HFD and control diet groups based on food intake at gestation, the relative abundance of *Bacteroides* in the gut microbiota was significantly depleted in neonates exposed to HFD during gestation (Chu et al., 2016).

Maternal diet is also critically involved in offspring neurodevelopment. Using a murine model, Ceasrine et al. (2022) reported that maternal HFD consumption from pre-pregnancy to gestation resulted in maternal lipid accumulation *in utero*, which disrupted 5-HT levels in the male fetal brain. They also reported that placental triglyceride accumulation was associated with pro-inflammatory signaling pathways in both sexes, and negatively correlated with brain 5-HT levels in males.

We previously examined the contribution of maternal gut microbiota composition at the end of lactation to offspring gut microbiota at weaning (3 weeks) and 14 weeks of age, comparing dams fed chow or a high-fat, high-sugar ‘cafeteria’ (Caf) diet (Hasebe et al., 2022). We found a strong influence of maternal gut microbiota on offspring gut microbiota composition at weaning; an influence that waned after weaning offspring onto either chow or Caf diets. To explore these associations in greater detail, in the present study we tracked changes in maternal gut microbiota at multiple times across pre-pregnancy, gestation and lactation to examine the dynamic effects of maternal obesogenic diet on microbiota across these stages. We sought to determine whether maternal microbiota composition is associated with offspring microbiota at two ages, weaning (3 weeks), proximal to maternal Caf diet exposure, and at 14 weeks old, after being weaned onto chow.

## Methods

### Ethics statement

The experimental protocol was approved by the Animal Care and Ethics Committee of the University of New South Wales (Ethics number:19/74A) in accordance with the guidelines for the use and care of animals for scientific purposes 8^th^ edition (National Health and Medical Research Council, Australia).

### Subjects

This study used a subset of animals from a larger study investigating the impact of maternal diet on offspring behaviour. Young adult female (approximately 7-8 weeks of age; body weight ∼200 g) and male (approximately 8-9 weeks of age; body weight ∼300 g) Sprague-Dawley rats were obtained from a commercial supplier (Animal Resource Centre, WA, Australia) and housed by sex 4 per cage in a colony room maintained at 18-22 degrees (12 h light/dark cycle). Water and standard chow (14 kJ/g, 65% carbohydrate, 22% protein and 13% fat; Premium Rat Maintenance diet, Gordon’s Stockfeeds, NSW, Australia) were continuously available. Following acclimatisation, female rats were weight-matched, then randomly allocated to receive chow (Chow; *n* = 10) or Caf (*n* = 15), a diet consisting of a selection of cakes, biscuits and protein sources e.g., meat pie and dim sims (food macronutrients in Supplemental Table 1). Foods varied daily, with chow and water always available, as described previously (Leigh et al., 2019).

The experimental design is shown in Figure 1A. After six weeks of diet, females were mated with chow fed males by co-housing two females and one male for five days. The male was then removed. Pregnancy was inferred based on weight gain and females were housed individually from approximately gestation day 16. Litters were standardised to six male and six female pups, where possible, on postnatal day (PND1). Dams and offspring were weighed every three days during lactation. From weaning (PND20) to 14 weeks of age, all pups were fed chow regardless of the maternal diet. At weaning and at 14 weeks of age, subsets of offspring (*n* = 1-2/sex/litter) were anesthetised by i.p. injection of ketamine/xylazine and decapitated. Retroperitoneal (RP) adipose tissue (identified as the triangular pad of fat attached to the lateral abdominal wall) was collected bilaterally.

**Figure 1.**
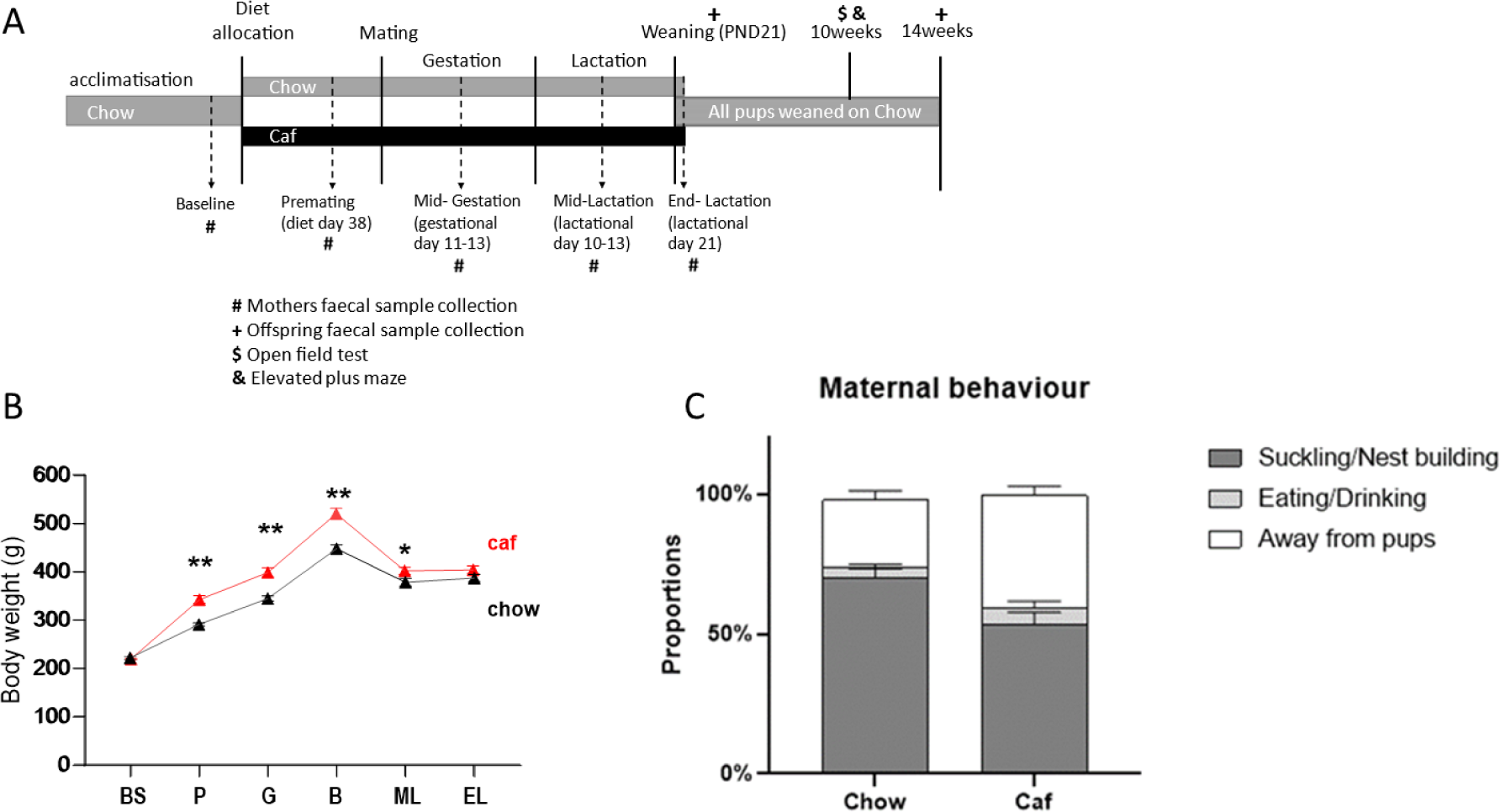
Experimental design **(A)**; Maternal body weight from baseline to end of lactation **(B)**, analysed by repeated measure ANOVA (time x diet). *p < .05, **p <.001, BS=baseline; P≈premating; G=gestation; B=birth; ML≈mid lactation; and EL≈eπd lactation.; and Maternal behaviour at lactation **(C)**, maternal behaviour during lactation was categorised as a) suckling/nest building, b) eating/drinking and c) time away from pups. Each proportion of maternal behaviour over total observations was calculated. Factorial ANOVA showed a significant effect of diet on maternal behaviour (F(_2(_69)=12.25, p < .001), with Caf dams spending less time suckling/nest building (p<.001) and more time away from pups than Chow dams (p <.Ol). Data expressed ± SEM n=10(chow) and 15 (Caf).

Maternal fecal samples were collected on five occasions (Figure 1A): at baseline (prior to introduction of Caf diet), pre-mating (diet day 38), mid-gestation (gestation day 11-13), mid-lactation (lactation day 10-13), and at the end of lactation when offspring were weaned (PND20). Offspring fecal samples were collected from the subset euthanised at weaning (PND20) and from their siblings continued on chow diet and euthanised at 14 weeks old.

### Behavioural measures

#### Maternal behaviour at lactation

During lactation, dams’ behaviour was recorded once daily (prior to feeding/weighing) and categorised as: suckling pups, eating, drinking, nest building, moving pups or other (away from pups and none of the other behaviours). The different categories of maternal behaviour were expressed as percentages for each dam and then analysed by factorial ANOVA. Endpoint Measures

When pups were weaned (PND20), dams were deeply anesthetised (ketamine/xylazine, i.p.) and body weight, girth, and nasoanal length measured. Blood was collected by cardiac puncture and rats were immediately decapitated. Retroperitoneal (RP) fat pads were weighed and snap-frozen. Plasma was stored at −30°C for analysis of leptin and insulin (CrystalChem Inc., Chicago, IL, USA) and triglyceride content (Roche triglyceride reagent, Sigma glycerol standard).

#### Faecal DNA extraction

Fresh faeces were collected, placed into a sterile tube and immediately frozen on dry ice. Faecal DNA extraction was performed using the PowerSoil® DNA Isolation Kit (Qiagen, Clayton, Victoria, Australia) according to the manufacturer’s instructions. DNA concentration and quality were measured using a DeNovix DS-11 Spectrophotometer (DeNovix, Inc., Delaware, USA) and stored at −80°C.

#### 16S rRNA gene amplicon sequencing and raw data analysis

Microbial community diversity was assessed by 16S rRNA gene amplicon sequencing (Illumina 2×250 bp MiSeq chemistry, V4 region, 515F-806R primer pair; Ramaciotti Centre for Genomics, UNSW Sydney) using a standard protocol (Caporaso et al., 2011). The sequence data were then analysed using Mothur (version 1.42.3, (Schloss et al., 2009)), which included removal of ambiguous bases and homopolymers longer than 15 base pairs, alignment with SILVA database version 132 (Quast et al., 2013), chimera checking with VSEARCH (version 2.13.3), and classification against the RDP Ribosomal Database training set (version18_03202018). Sequences were clustered into operational taxonomic units (OTU) at 97% nucleotide identity to generate an OTU count table and a taxonomic classification file. Commands were derived from MiSeq SOP (Kozich et al., 2013) and modified as required. Sequence data were subsampled to n = 4191 total clean reads/sample.

#### Statistical analysis

Data were analysed using SPSS (v28, IBM). Effects of the diet on maternal body weight during pregnancy were assessed in a mixed-ANOVA with factors of maternal diet (Chow or Caf) and time (baseline, premating, mid-gestation, mid-lactation and end-lactation), applying a Greenhouse-Geisser correction where appropriate. Dams’ endpoint measures were assessed by independent samples t-tests. Endpoint measures in weanlings and at 14 weeks were assessed in two-way ANOVAs with maternal diet (Chow or Caf) and sex (male or female) as between-subjects factors. OTU tables were standardised by dividing feature read counts by total number of reads in each sample to calculate relative abundances. Standardised data were then square root transformed and sample resemblances were calculated using Bray-Curtis similarities. β diversity was assessed by Non-metric Multi-dimensional Scaling (NMDS) plots, Permutational Multivariate Analysis of Variance (PERMANOVA), and Permutational Analysis of Multivariate Dispersions (PERMDISP) with Bray-Curtis resemblance matrices. Analyses were completed using PRIMER-e v7 (Primer-e Ltd., Plymouth, UK)(Clarke, 1993). Repeated measures ANOVA was used for analyses of relative abundance in dams with significance level set *p* <0.025 and the Bonferroni test was used to control for inflation of the Type 1 error rate by multiple comparisons. SourceTracker (Knights et al., 2011) assessed similarities between maternal biota at multiple time points and offspring microbiota via the Galaxy web application (Feng et al., 2017; Jalili et al., 2020). Figures were generated in GraphPad Prism v9 and PRIMER-e v7. Results are expressed as mean ± SEM and considered significant at p<0.05.

## Results

### Maternal behaviour during lactation and endpoint measures

Figure 1A shows the experimental design in this study. Analysis of maternal body weight from baseline to lactation revealed a significant time x diet interaction (F_(2.86,62.84)_=17.11, *p*<.001), and main effects of time (F_(2.86, 62.84)_=71.07, *p*<.001) and diet (F_(1,22)_=15.11, *p*<.001), indicating that the Caf diet significantly increased body weight at premating, mid-gestation, and mid-lactation periods (Figure 1B). As shown in Table 1, gestational weight gain did not differ significantly between Chow and Caf dams, but during lactation body weight declined in the Caf group while remaining stable in the Chow group (*p*<.001, Table 1). Consequently, at the end of lactation, body weight did not differ between groups. Nonetheless, RP fat mass, levels of blood glucose and plasma leptin were higher in Caf than Chow dams. Maternal behaviour (suckling/nest building; eating/drinking, and away from pups) during lactation differed between diet groups (F_(2,69)_=12.25, *p*<.001). Compared with Chow dams, Caf dams spent significantly less time suckling/nest building (*p*<.001), and less time attending to pups (*p*<.01, Figure 1C).

### Maternal diet effects on microbiota composition across pregnancy

Repeated measures ANOVA indicated a significant interaction between time and Caf diet (F_(4,60)_=8.50, *p*<.001) and a main effect of time (F_(4,60)_=3.60, *p*<.05) on richness (Figure 2A), which was significantly higher in Caf than Chow dams at mid- and end-lactation (*p*<.001 and *p*<.01 respectively). Similarly, a significant interaction between time and Caf diet was observed for Shannon index (F_(4,60)_=3.02, *p*<.05), which was significantly higher in Caf dams at mid- and end-lactation compared with Chow dams (*p*<.05 and *p*<.01 respectively) (Figure 2B). In a principal coordinate analysis (PCO) with Bray-Curtis resemblance (Figure 2C), PCO1 axis represents a variation on diet and PCO2 axis represents progression of pregnancy. For Chow dams, permutational MANOVA (PERMANOVA) indicated a significant main effect of time on microbiota composition (Pseudo-F = 3.51, *p*<.01). Each timepoint differed significantly (*p* values ranging from 0.0002 to 0.0048), except premating and gestation timepoints (*p*=.0525, Supplemental Table 2.1). However, analyses of centroid distance (Distance-based test for homogeneity of multivariate dispersions, PERMDISP) showed a significant difference between time points (F_(4, 35)_ = 5.87, *p*<.01), including between baseline and mid-lactation (*p*<.01) and end-lactation (*p* <.001), and between premating and end of lactation (*p*<.05, Supplemental Table 2.2). Hence, differences in dispersions may contribute to the effects of PERMANOVA in Chow dams. To further investigate the dispersion effects, we conducted Analysis of Similarity (ANOSIM) to assess similarities between each timepoint (overall R statistic R=0.55, *p*=0.0001), with significant pairwise comparisons between all times (R ranging from 0.227 to 0.864; *p* values 0.043 to 0.0002). For Caf dams, PERMANOVA revealed significant differences in β diversity across time (Pseudo-F=3.56, *p*<.001), with all time points significantly different from one another (Supplemental Table 2.1). Unlike Chow dams, however, no differences in dispersion were observed (PERMDISP: F_(4,25)_ = 0.51, *p*>.05, see Supplemental Table 2.2).

**Figure 2.**
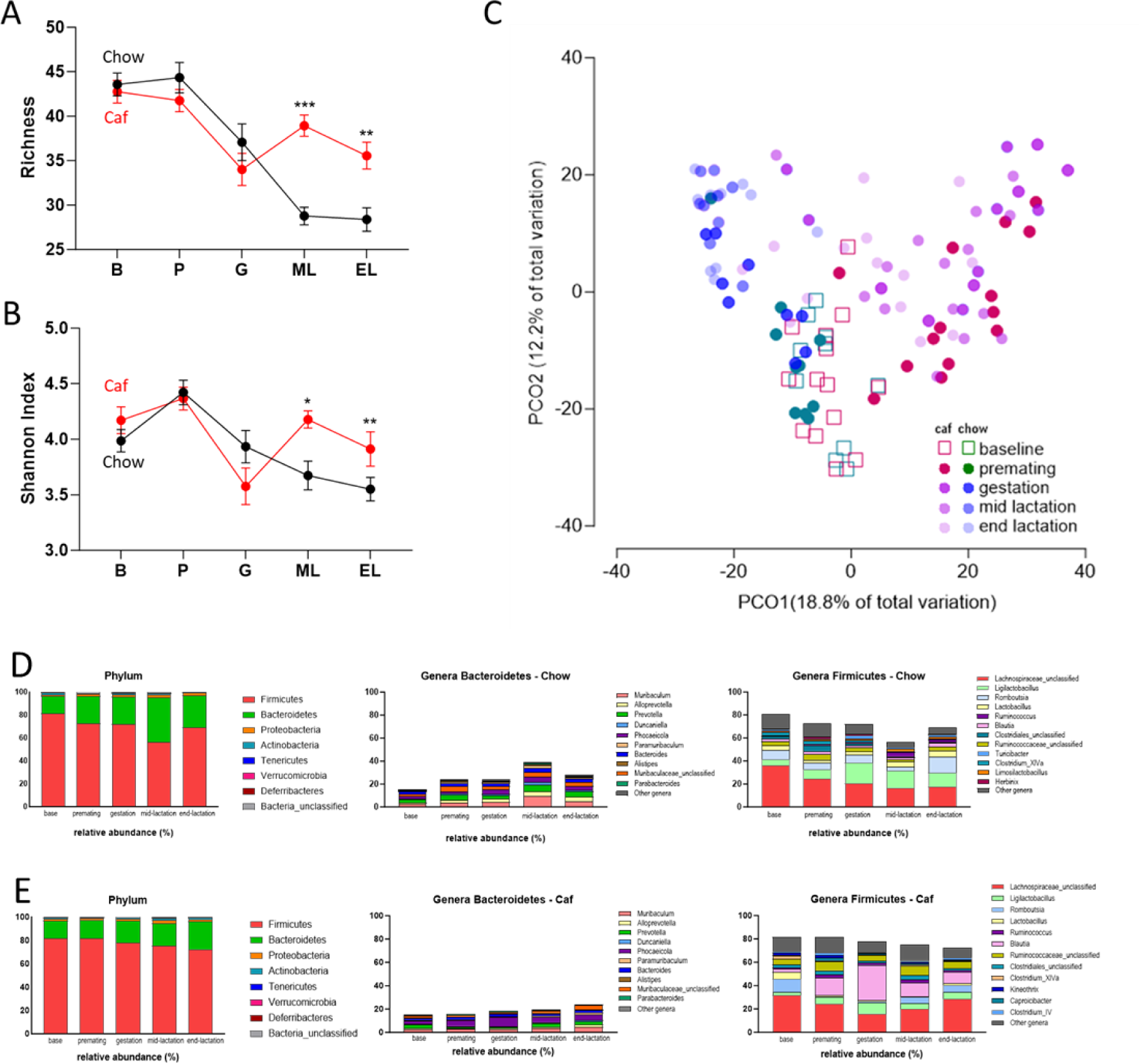
Gut microbiome development with progression of pregnancy and effect of Cafeteria diet Gut microbiota of mothers fed chow diet were assessed at multiple timepoints, baseline (B), premating (P), gestation (G), lactation (ML) and end of lactation (EL). Alpha diversity - Richness **(A)**; and Shannon index **(B)** Different letters indicate statistically significance by Bonferroni post hoc comparison, *p <05, **p<.01,***p<.001 compared with Chow dams. Data are displayed ± SEM; Principal coordinate analyses (PCO) following square root transformation and Bray-Curtis resemblance of relative abundance data at the OTU level in Chow and Caf dams {C}. Representative taxa of chow (DĮ and Caf **(E)** fed dams during pregnancy at phylum level, and for genera Bacteroidetes and genera Firmicutes.

The change in *Bacteroidetes* abundance across time differed between groups (F_(4,60)_=5.70, *p*<.001) with significantly higher *Bacteroidetes* abundance in Chow than Caf dams at gestation and mid-lactation (*p*<.05 and *p*<.001 respectively). *Firmicutes* abundance also differed between groups over time (F_(4,60)_=5.29, *p*<.01) with significantly lower *Firmicutes* in Chow than Caf dams at gestation and mid-lactation (*p*<.05 and *p*<.001 respectively). Within group analyses in Chow dam indicated significant differences in abundance of *Bacteroidetes* (F_(4,24)_=3.76, p<.05) and *Firmicutes* (F_(4,24)_=5.03, *p*<.01) from baseline to end lactation (Figure 2D). *Bacteroidetes* abundance was significantly higher at lactation than baseline (*p*<.05) while *Firmicutes* abundance was significantly lower at lactation than baseline and premating (both *p*s<.05). In the case of Caf dams, repeated measure ANOVA indicated that there was no significant difference in *Bacteroidetes* and *Firmicutes* abundance levels from baseline to end lactation (Figure 2E).

### Endpoint measures in male and female offspring at weaning and 14 weeks of age

Table 3 summarises endpoint measures in offspring at weaning and 14 weeks of age. Factorial ANOVA with maternal diet (Chow versus Caf) and sex (male versus female) confirmed that maternal Caf diet consumption significantly increased weanling blood glucose (F_(1,44)_=15.771, *p*<.001), plasma leptin (F_(1,44)_=35.059, *p*<.001) and retroperitoneal (RP) fat mass (F_(1,44)_=46.994, *p*<.001). It also confirmed that male weanlings had significantly greater RP fat mass than females (F_(1,44)_=4.411, *p*<.05). There were no significant effects of maternal diet or sex on weanling body weight.

**Table 1.**
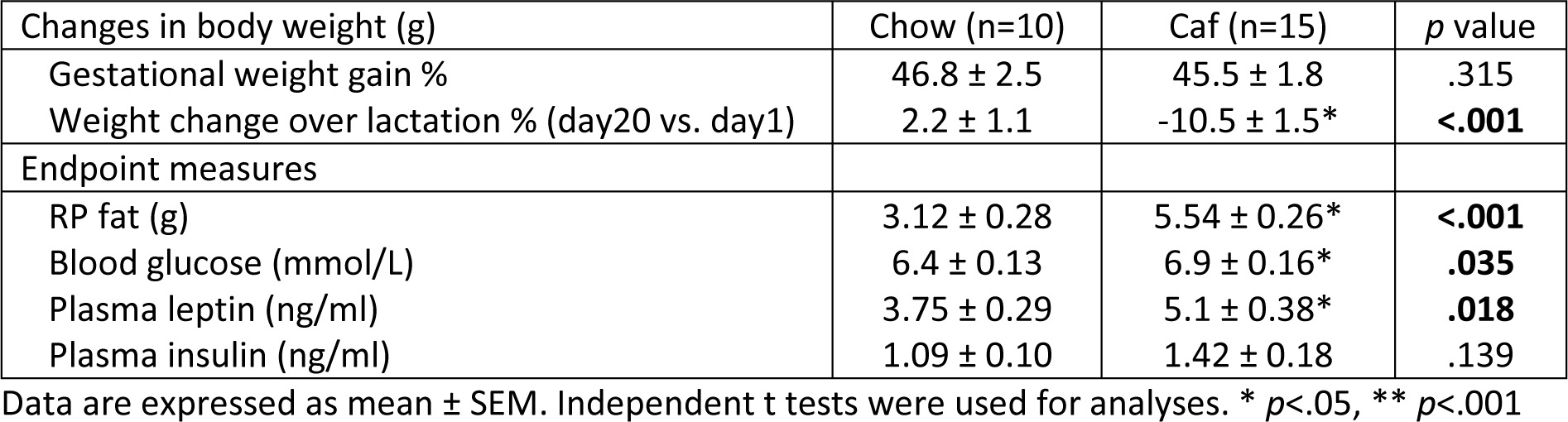
Metabolic parameter in dams fed chow or Caf diet from pre-pregnancy to weaning.

**Table 3.**
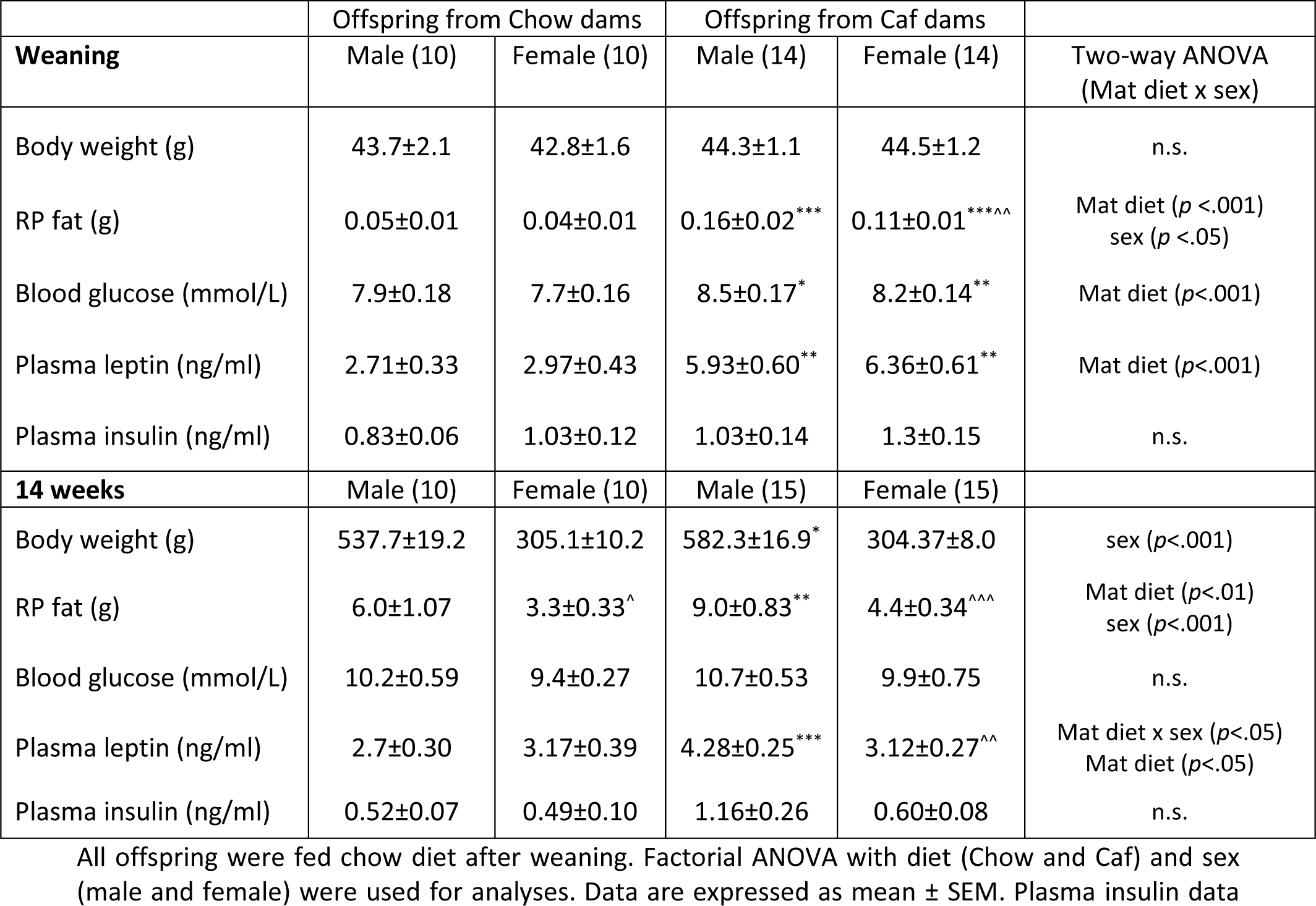
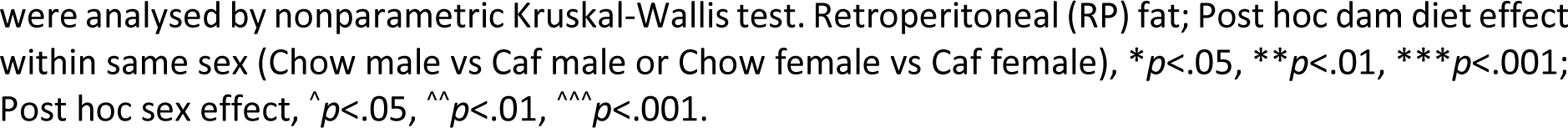
Offspring endpoint measures – weaning and 14 weeks of age.

At 14 weeks of age, maternal Caf diet was associated with greater RP fat mass (F_(1,46)_=8.466, *p*<.01), and male offspring from Caf dams were significantly heavier than those from Chow dams (*p*<.05). There was a significant interaction between sex and maternal diet (F_(1,46)_=7.202, *p*<.05) and a significant maternal diet effect (F_(1,46)_=6.337, *p*<.05) on plasma leptin levels. Bonferroni post hoc comparisons confirmed that maternal Caf diet significantly elevated plasma leptin in male but not in female offspring (*p*>.05; Table 3).

### Influence of maternal microbiota on offspring microbiota

Offspring α diversity was analysed by two-way ANOVA with factors of age (weanling versus 14 weeks) and maternal diet (Chow versus Caf). Species richness and Shannon index significantly increased with age (Species richness F_(1,94)_=44.30, *p*<.0001; Shannon index F_(1,94)_=221.0, *p*<.0001) in offspring from both Chow and Caf dams (*p*s <.001 to <.0001; Figure 3A-B). There was no significant interaction between maternal diet and time, and no main effect of maternal diet.

**Figure 3.**
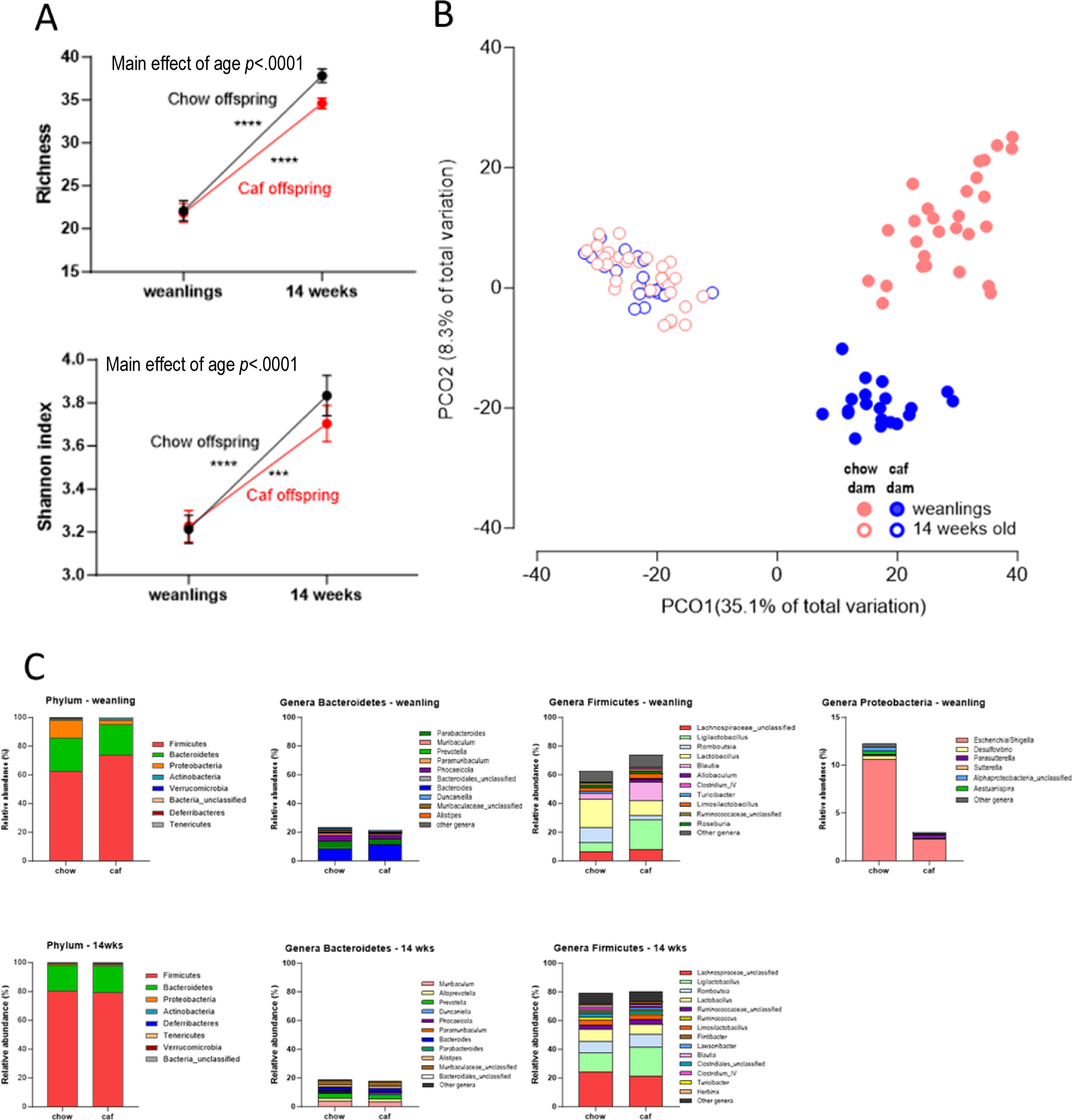
Effect of maternal diet on offspring microbiota development (weaning to 14 weeks, all offspring fed Chow after weaning). Alpha diversity - Richness and Shannon index **(A)**. Data are displayed as mean ± SEM. Weanlings n=20-28; 14 weeks n=20-30. Statistical significance by Bonferroni post hoc comparison, * p<.05; *** p <.001; ****p<,0001. Principal coordinate analyses (PCO) following square root transformation and Bray-Curtis resemblance of relative abundance data at the OTU level in weanlings and 14 weeks offspring **(B)**. Relative abundance in weanlings and 14 weeks offspring **(C)**.

Figure 3B shows PCO of weanlings and 14 week-old offspring. PCO1 axis shows a separation by age (weanling or 14 weeks), and PCO2 axis shows a separation by maternal diet in weanling, but not in 14 week-old offspring. All weanlings and 14 weeks offspring were fed chow after weaning. When Chow and Caf offspring β diversities were directly compared at weaning and 14 weeks of age, maternal Caf diet had a significant effect on β diversity in Caf versus Chow weanlings (PERMANOVA, Pseudo F=10.18, *p*=0.0001; PERMDISP F=2.74, *p*=0.1), and 14-week-old offspring (PERMANOVA, Pseudo F=2.4679, *p*=0.005; PERMDISP F=1.23, *p*= 0.38). Sex did not differentiate β diversity at either age and the maternal diet effect did not interact with sex. Descriptive representative taxa in weanlings and 14-week-old offspring are shown in Figure 3C. It appears that maternal Caf diet consumption virtually eliminated *Escherichia* and instead, promoted *Firmicutes* abundance in weanlings.

### Similarity of maternal and offspring gut microbiota

SourceTracker analysis examined whether the microbiota of dams were related to their offspring gut microbiota composition (Figure 4). Chow weanling microbiota most closely resembled the maternal microbiota at mid- and end-lactation (blue and purple respectively). In contrast, Caf weanling microbiota most closely resembled the microbiota at mid-gestation. In adult offspring from Chow dams, microbiota composition more closely resembled the microbiota at gestation (green) along with contributions from lactation microbiota, whereas microbiota of adult offspring from Caf dams resembled the baseline and end-lactation maternal microbiota, perhaps due to them being weaned onto chow and continuing on this diet from 3 to 14 weeks of age.

**Figure 4.**
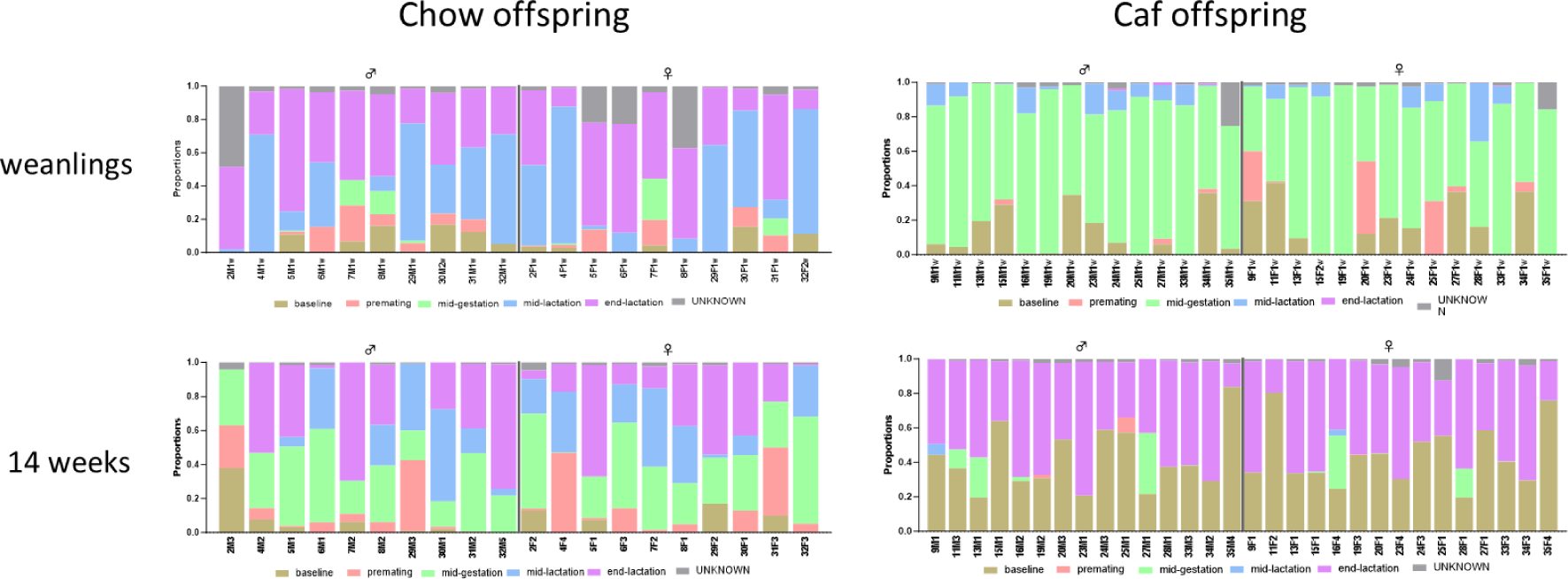
Influence of maternal microbiota on offspring at weaning and 14 weeks. Source Tracker analysis shows influence of gut microbiota in Chow fed dams on weaners and 14-week offspring consuming chow (left panel); and Caf fed dams on weaners and 14-week old offspring consuming chow (right panel). Each bar represents an individual animal, including male, female sibling pairs from 10 different dams, as numbered. Mothers’’ gut microbiota at baseline, premating, gestation, lactation and end of lactation were computed as source in order to evaluate the proportion of the microbiota at each time point that contributed to her offspring gut microbiota at weaning and 14 weeks of age.

## Discussion

This experiment showed that consumption of a high-fat, high-sugar, cafeteria diet altered the trajectory of gut microbiota composition of rat dams across pregnancy and lactation. The decrease in microbiota species richness and Shannon index observed in Chow dams during lactation was absent in Caf dams. Similarity, the fluctuations in two major phyla, Bacteroidetes and Firmicutes across gestation and lactation in Chow dams were absent in Caf-fed dams. Identifying the impact of diet on gut microbiota composition across human pregnancy can be obscured by numerous confounding factors such as intake of supplements, and differences in overall dietary intake. Hence, the present results might shed light on the diverse findings from studies on maternal microbiota in humans.

Although PERMANOVA indicated that both Chow and Caf maternal gut microbiota differed from baseline to lactation, PCO and PERMDISP suggested that Chow maternal gut microbiota shows more distinct change between each time point whereas Caf maternal gut microbiota differed less over time, possibly due to disruptive effects of Caf diet on the gut microbiota. Previous non-human studies have reported changes in maternal gut microbiota across pregnancy (Ji et al., 2019; Paul et al., 2018; Zhang et al., 2023), which in line with the changes in Chow dams in our study. In contrast, the present study found that Caf diet induced moderate changes in maternal gut microbiota during pregnancy, which may be one of the factors that contribute to the diverse findings in human pregnancy (DiGiulio et al., 2015; Rasmussen et al., 2020; Yang et al., 2020).

The present experiment also found that maternal Caf diet suppressed changes in *Bacteroidetes* and *Firmicutes* abundance during pregnancy and lactation in comparison to their abundance in dams fed chow. More specifically, the changes in the two phyla observed here were an interaction between Caf diet and the two major phyla with the progression of pregnancy, rather than the Firmicutes/Bacteroidetes ratio, an indicator of health or disease states (Magne et al., 2020). Whether this Caf diet induced suppression of changes in the two major phyla influences outcome of pregnancy, offspring gut microbiota, and offspring development may require further investigation.

Maternal Caf diet consumption generated a robust obesity phenotype in dams, with a 20% difference in body weight prior to mating persisting to parturition. Of note, Caf dams lost weight during lactation such that dam groups did not differ in weight at endpoint, though adiposity and plasma leptin remained higher in Caf dams, indicating a residual metabolic impairment. Offspring born to Caf dams exhibited a mild obesity phenotype, with greater adiposity, plasma leptin and blood glucose than Chow dam offspring at weaning, and increased adiposity in adulthood, with a sex-specific increase in body weight in males, consistent with our previous work (Tajaddini et al., 2022).

Caf dams tended to their offspring less during lactation, suggesting maternal care was compromised. Caf diet consumption affects reward systems which subsequently may alter social behaviours, including maternal care, in rodents (Lalanza and Snoeren, 2021). For instance, eight days of Caf diet exposure during lactation in rats resulted in more time spent licking and grooming the pups (Speight et al., 2017). Similarly, Ribeiro et al. (2018) reported in mice that exposure to Caf diet during gestation increased maternal nursing, nest building, and licking of pups during lactation. In contrast, Caf dams in our study spent less time on nursing behaviour. The discrepancy in maternal behaviour may be due to the duration and timing of Caf diet exposure, as both studies by Speight et al. (2017) and Ribeiro et al. (2018), involved only brief Caf diet exposure during lactation (Speight et al., 2017) and gestation (Ribeiro et al., 2018), whereas the present study involved additional exposure prior to mating. Maternal Caf diet consumption altered the offspring gut microbiota, despite offspring being weaned onto chow diet. Maternal Caf diet significantly changed β diversity but not α diversity in their offspring at weaning. The degree of similarity between the maternal and offspring gut microbiota at the different timepoints differed for Chow and Caf weanlings. Although both were fed chow post weaning, differences in phenotype between Chow and Caf offspring were still apparent at 14 weeks of age. Our previous study showed that adult offspring gut microbiota were more affected by post weaning diet (chow vs. Caf), than by maternal diet (Hasebe et al. 2020). SourceTracker analysis show more nuanced maternal influences on offspring gut microbiota which were not evident through α- and β diversity measures, instead showing different patterns of contribution of maternal gut microbiota across the extremes of diet.

### Limitations

Our study showed alterations of maternal gut microbiota during pregnancy to lactation under the influence of a cafeteria diet that mimics high-sugar, high-fat and highly processed diets eaten by many people, including pregnant mothers, around the world. While the diet regimen in this rat study has strong relevance to human dietary patterns, our focus was on the impact of diet on gut microbiota, and hence, we minimised potential confounding factors in our laboratory setting, which is not feasible in human settings. This may limit the applicability of our findings for humans. Moreover in this study we were unable to assess the fibre content of each diet (Chow and Caf) along with other food additives which could interact with macronutrients and subsequently differentially influence the gut microbiota, between chow and Caf fed animals (Morrison et al., 2020).

## Conclusion

This study showed that long term obesogenic diet consumption alters the trajectory of maternal gut microbiota from premating to lactation. Maternal Caf diet programs offspring metabolic development, but also disrupts connections between maternal gut microbiota and offspring gut microbiota, and as such, it can impact on the development of offspring gut microbiota in later life. While the gut microbiota of Caf offspring differed from that of Chow offspring at weaning, the difference was less marked at 14 weeks of age when offspring were consuming chow.

## Acknowledgement and right retention policy statement

This research was funded in whole or part by the National Health and Medical Research Council to MJM and RFW [project grant number: APP1161418]. For the purposes of open access, the author has applied a CC BY public copyright license to any Author Accepted Manuscript version arising from this submission.

## Data availability

The sequence data for this study will be available upon request.

## List of Supplemental data

**Supplemental table 1.**
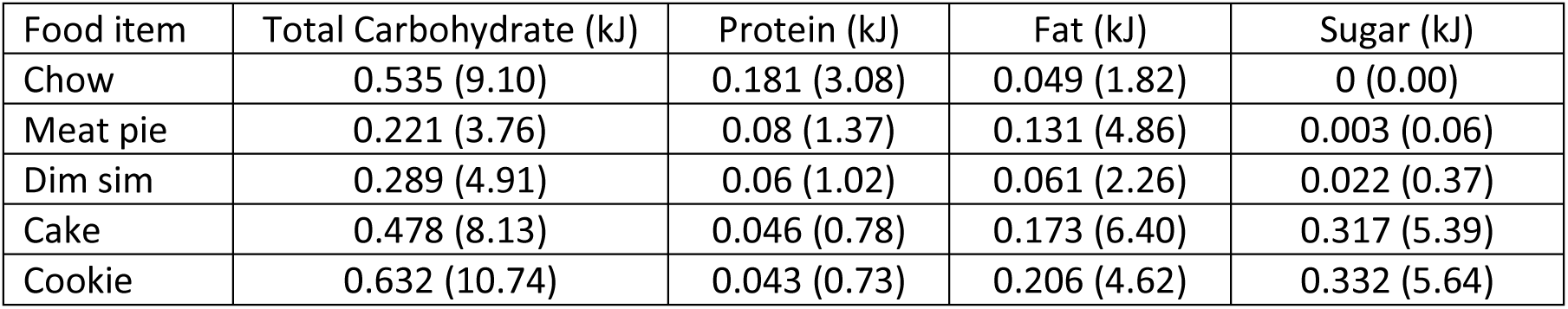
Macronutrients (mg/g per each item)

**Supplemental Table 2.1.**
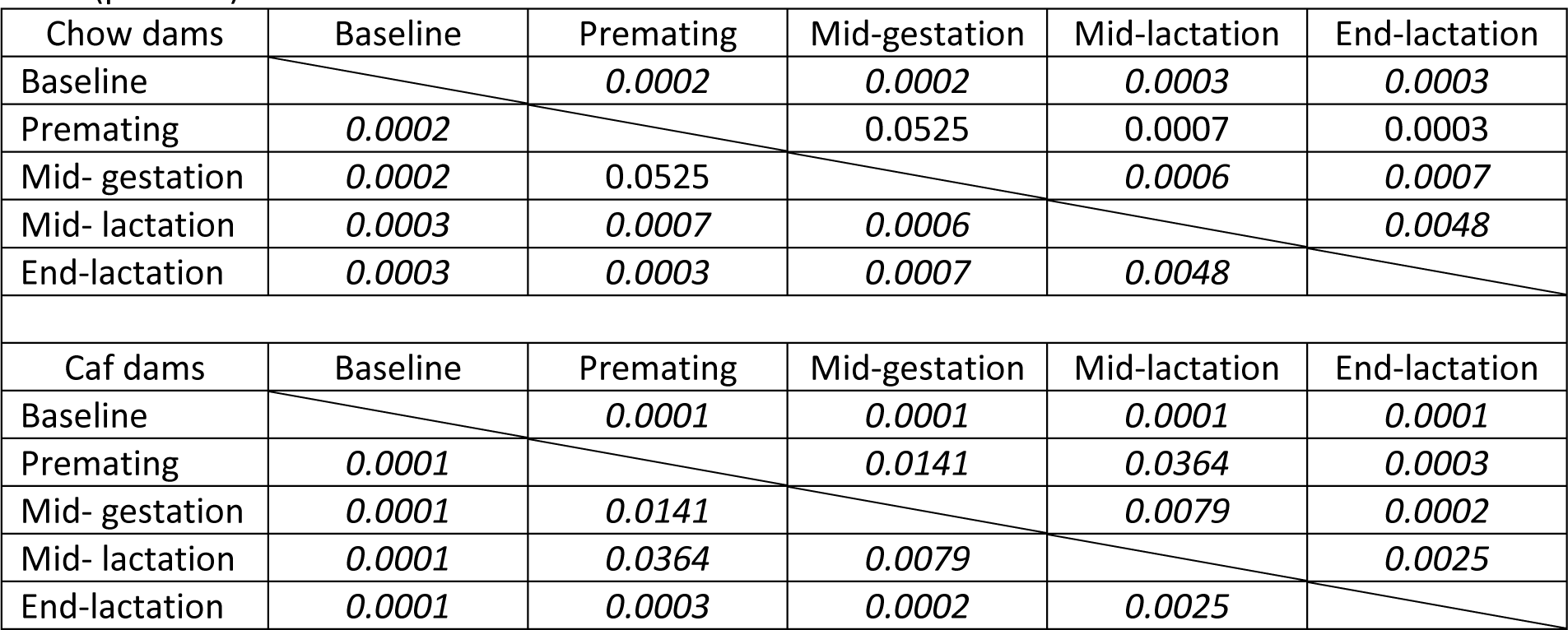
Summary of PERMANOVA pairwise comparisons between time points in dams (p values)

**Supplemental Table 2.2.**
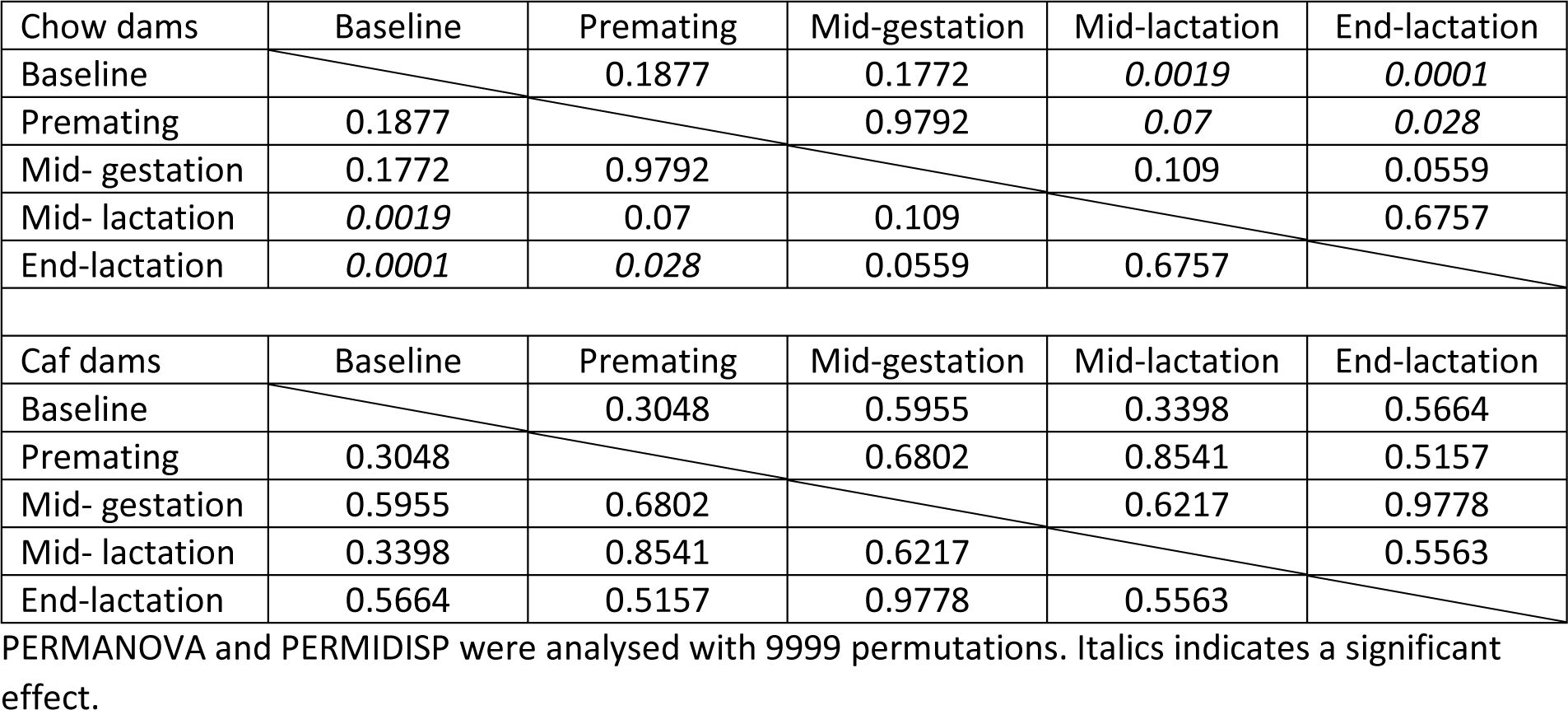
Summary PERMIDISP pairwise comparisons between time points in dams (p values)

