## Supplemental tables for "Impact of maternal obesogenic diet on maternal and offspring microbiome development"

List of Supplemental data

Supplemental Table 1 – Macronutrients per food item comprising Cafeteria diet
Supplemental Table 2.1 – Summary of PERMANOVA pairwise comparisons between time points in dams (p values)
Supplemental Table 2.2 – Summary PERMIDISP pairwise comparisons between time points in dams (p values)

**Supplemental table 1 Macronutrients (mg/g per each item)**

| Food item | Total Carbohydrate (kJ) | Protein (kJ) | Fat (kJ) | Sugar (kJ) |
|---|---|---|---|---|
| Chow | 0.535 (9.10) | 0.181 (3.08) | 0.049 (1.82) | 0 (0.00) |
| Meat pie | 0.221 (3.76) | 0.08 (1.37) | 0.131 (4.86) | 0.003 (0.06) |
| Dim sim | 0.289 (4.91) | 0.06 (1.02) | 0.061 (2.26) | 0.022 (0.37) |
| Cake | 0.478 (8.13) | 0.046 (0.78) | 0.173 (6.40) | 0.317 (5.39) |
| Cookie | 0.632 (10.74) | 0.043 (0.73) | 0.206 (4.62) | 0.332 (5.64) |

**Supplemental Table 2.1** Summary of PERMANOVA pairwise comparisons between time points in dams (p values)

| Chow dams | Baseline | Premating | Mid-gestation | Mid-lactation | End-lactation |
|---|---|---|---|---|---|
| Baseline | | *0.0002* | *0.0002* | *0.0003* | *0.0003* |
| Premating | *0.0002* | | 0.0525 | 0.0007 | 0.0003 |
| Mid- gestation | *0.0002* | 0.0525 | | *0.0006* | *0.0007* |
| Mid- lactation | *0.0003* | *0.0007* | *0.0006* | | *0.0048* |
| End-lactation | *0.0003* | *0.0003* | *0.0007* | *0.0048* | |

| Caf dams | Baseline | Premating | Mid-gestation | Mid-lactation | End-lactation |
|---|---|---|---|---|---|
| Baseline | | *0.0001* | *0.0001* | *0.0001* | *0.0001* |
| Premating | *0.0001* | | *0.0141* | *0.0364* | *0.0003* |
| Mid- gestation | *0.0001* | *0.0141* | | *0.0079* | *0.0002* |
| Mid- lactation | *0.0001* | *0.0364* | *0.0079* | | *0.0025* |
| End-lactation | *0.0001* | *0.0003* | *0.0002* | *0.0025* | |

**Supplemental Table 2.2** Summary PERMIDISP pairwise comparisons between time points in dams (p values)

| Chow dams | Baseline | Premating | Mid-gestation | Mid-lactation | End-lactation |
|---|---|---|---|---|---|
| Baseline | | 0.1877 | 0.1772 | *0.0019* | *0.0001* |
| Premating | 0.1877 | | 0.9792 | *0.07* | *0.028* |
| Mid- gestation | 0.1772 | 0.9792 | | 0.109 | 0.0559 |
| Mid- lactation | *0.0019* | 0.07 | 0.109 | | 0.6757 |
| End-lactation | *0.0001* | *0.028* | 0.0559 | 0.6757 | |

| Caf dams | Baseline | Premating | Mid-gestation | Mid-lactation | End-lactation |
|---|---|---|---|---|---|
| Baseline | | 0.3048 | 0.5955 | 0.3398 | 0.5664 |
| Premating | 0.3048 | | 0.6802 | 0.8541 | 0.5157 |
| Mid- gestation | 0.5955 | 0.6802 | | 0.6217 | 0.9778 |
| Mid- lactation | 0.3398 | 0.8541 | 0.6217 | | 0.5563 |
| End-lactation | 0.5664 | 0.5157 | 0.9778 | 0.5563 | |

PERMANOVA and PERMIDISP were analysed with 9999 permutations. Italics indicates a significant effect.
